# Characterization of a vaccine-elicited human antibody with sequence homology to VRC01-class antibodies that binds the C1C2 gp120 domain

**DOI:** 10.1101/2021.08.21.457217

**Authors:** Matthew D. Gray, Junli Feng, Connor E. Weidle, Kristen W. Cohen, Lamar Ballweber-Fleming, Anna J. MacCamy, Crystal N. Huynh, Josephine J. Trichka, David Montefiori, Guido Ferrari, Marie Pancera, M. Juliana McElrath, Leonidas Stamatatos

## Abstract

Broadly HIV-1 neutralizing VRC01-class antibodies bind the CD4-binding site of the HIV-1 envelope (Env) and contain VH1-2*02-derived heavy chains paired with light chains expressing five amino acid long CDRL3s. Their unmutated forms do not recognize Env or neutralize HIV-1. The lack of elicitation of VRC01-class antibodies in human clinical trials could potentially be due to the absence of activation of the corresponding naïve B cells by the vaccine Env immunogens. To address this point directly, we examined Env-specific BCR sequences from participants in the HVTN 100 clinical trial. Of all the sequences analyzed only one displayed sequence homology to VRC01-class antibodies, but the corresponding antibody (FH1) recognized the C1C2 gp120 domain. For FH1 to switch epitope recognition to the CD4-binding site, alterations in both the CDRH3 and CDRL3 were necessary. Our findings support the use of specifically designed immunogens to activate VRC01-class B cells in future human vaccine trials.

## INTRODUCTION

HIV-1 broadly neutralizing antibodies (bnAbs) target diverse areas of the viral Env spike, including the CD4-binding site (CD4-BS), the apex region, the N332 glycan patch, the interface between the gp120 and gp41 subunits and the gp41 subunit itself (Burton and Hangartner, 2016; Kwong and Mascola, 2012; McCoy and Burton, 2017). bnAbs that bind the same Env region and share common genetic and/or structural features are grouped into ‘classes’ (Kwong and Mascola, 2012). The anti-CD4-BS VRC01-class antibodies are among the most potent bnAbs known (Sok and Burton, 2016), they prevent SHIV-infection of rhesus macaques and HIV-1-infection of humanized mice (Balazs et al., 2012; Shingai et al., 2014) and are targets for HIV-1 vaccine development.

VRC01-class antibodies have been isolated from several HIV-1-infected subjects, but their heavy chain (HC) variable domains are derived exclusively from the VH1-2*02 gene allele and are paired with light chains (LCs) expressing five amino acid long CDRL3s (Huang et al., 2016; Sajadi et al., 2018; Scheid et al., 2011; Umotoy et al., 2019; Wu et al., 2015; Wu et al., 2011; Zhou et al., 2010; Zhou et al., 2013). Their VH and VL genes are extensively mutated (Scheid et al., 2011; Wu et al., 2011; Zhou et al., 2013). However, despite extensive amino acid sequence diversity amongst them, VRC01-class antibodies adopt similar overall structures and engage the CD4-BS with similar angles of approach (Zhou et al., 2013).

In contrast to their mature mutated forms, recombinant antibodies generated from their inferred unmutated antibody sequences (‘germline’, gl) do not bind recEnv or neutralize HIV-1 (Jardine et al., 2013; McGuire et al., 2014b; McGuire et al., 2013; Zhou et al., 2010). These observations lead to the proposal that Envs employed as immunogens in human clinical trials, would not have activated naïve B cells expressing glVRC01-class BCRs (Jardine et al., 2013; McGuire et al., 2013; Stamatatos et al., 2017; Zhou et al., 2010). If true, this would in part explain the absence of VRC01-like neutralizing activities in sera collected from vaccinees. However, experimental evidence for this is presently lacking.

Participants of the HVTN100 phase 1-2 clinical trial were immunized with ALVAC vectors (vCP2438), expressing clade C ZM96 gp120 with the gp41 transmembrane sequence of the subtype B LAI strain and the *gag* and *protease* of LAI, at months 0 and 2 and then co-administered with recombinant clade C Env gp120 proteins (TV1.C and 1086.c) formulated in squalene-based adjuvant MF59 at 4, 6 and 12 months (Bekker et al., 2018; Shen et al., 2020). Known glVRC01-class antibodies do not bind these Env proteins and thus, they were not expected to have activated B cells expressing VRC01-class BCRs *in vivo*. Following an in-depth analysis of BCRs expressed by vaccine-specific gp120+ memory B cells isolated at the end of the immunization regimen from a subset of vaccinated subjects, 339 BCRs were analyzed. Only one, FH1, displayed classic VRC01-like HC and LC characteristics, i.e., a HC derived from VH1-2*02 paired with a k3-20 LC expressing a 5 amino acid long CDRL3. The amino acid sequence homology between FH1 and germline VRC01 (outside the CDRH3 and CDRL3) is 95% (95/100) in the HC and 98% (92/94) in the LC. The isolation of FH1 potentially suggests that Env-based immunogens that are not specifically designed to activate VRC01-class B cells may do so, albeit infrequently.

Here, we characterized the structural, antigenic and functional properties of FH1 and determined that, in contrast to VRC01-class antibodies, it recognizes the C1C2 domain of gp120, where non-neutralizing antibodies with ADCC activities also bind (Acharya et al., 2014; Ferrari et al., 2011). We also report that for FH1 to switch its epitope-specificity towards the CD4-BS, both CDRH3 and CDRL3 domains had to be modified to those of VRC01. By engineering B cells to express either FH1 or glVRC01 BCRs we were able to confirm that the latter can only be activated by specifically designed germline-targeting immunogens, but not by the Env used in the HVTN100 trial. As such, our study provides evidence in support of the clinical evaluation of Env immunogens that have been specifically designed to target VRC01-expressing B cells.

## RESULTS

### Isolation of an antibody with VRC01-like genetic characteristics

HVTN 100 was a phase 1/2 clinical trial of a RV144-like vaccine regimen which had been modified with subtype C specific immunogens for the South African epidemic (Fig S1a) (Bekker et al., 2018). Two weeks after the fifth immunization, vaccine-specific 1086.C gp120+ IgG+ and vaccine-nonspecific 1086.C gp120-IgG+ B cell populations were isolated and their VH/VL genes sequenced from 14 vaccine recipients.

Env-specific B cells were single-cell sorted from the vaccine recipients with the highest proportion of VH1-2*02 gene usage. Of the Env-specific B cells from which we recovered paired VH and VL genes, we identified one VH1-2*02 HC was paired with a Vκ3-20 with a 5 amino acid-long CDRL3 (“FH1”; Fig. 1a).

**Figure 1.**
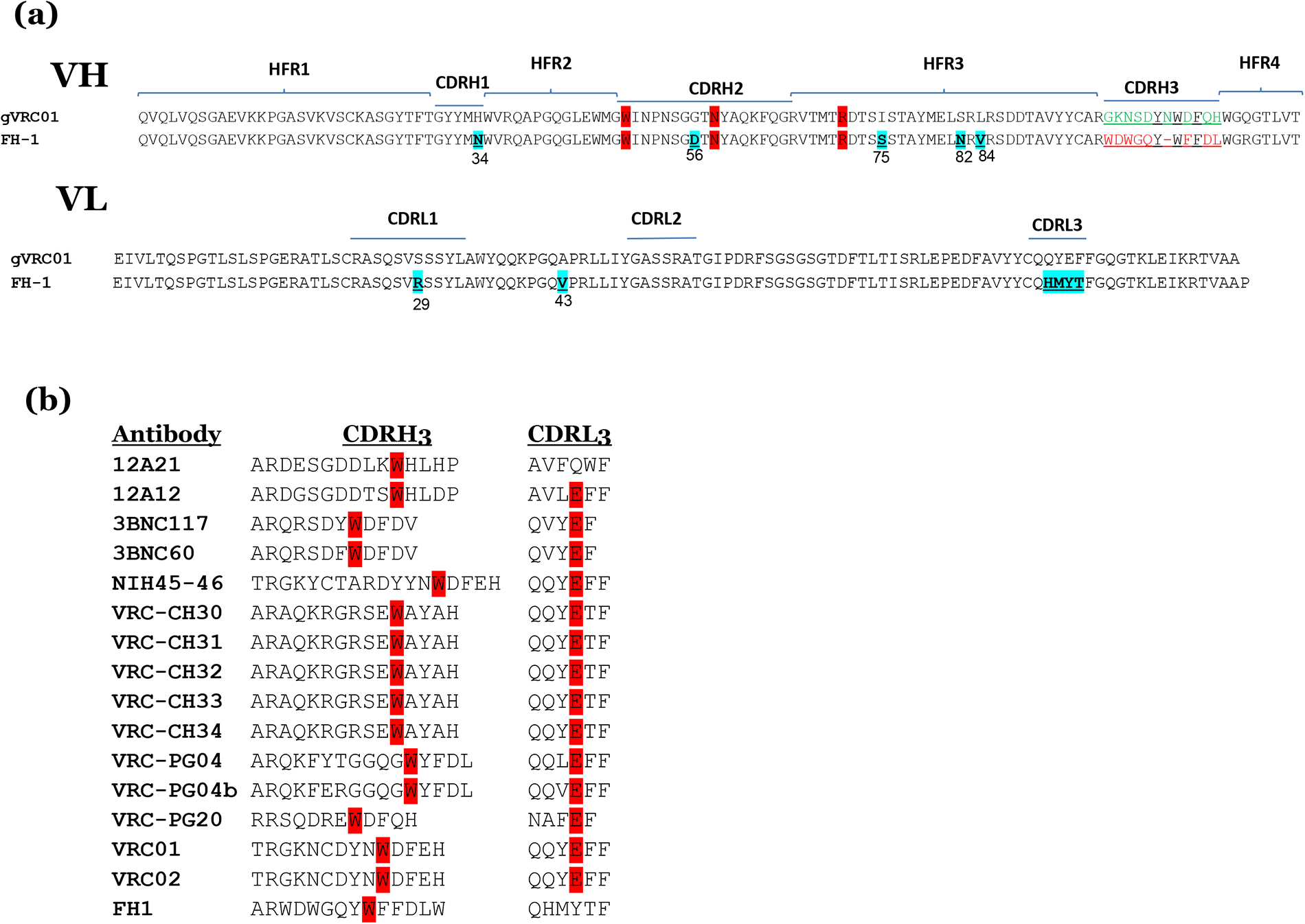
Amino acid sequence of FH1 and VRC-1-class antibodies. (**a**) Alignment of the VH and VL amino acid sequences of FH1 and glVRC01. Amino acids that differ between the two antibody sequences are highlighted in cyan and are underlined. The three gene-encoded amino acids in the VH1-2*02 domains (Trp50_HC_, Asn58_HC_ and Arg71_HC_, Kabat numbering) that play key roles in the interaction between VRC01-class antibodies and the CD4-BS of Env are highlighted in red. (**b**) Sequence alignment of the CDRH3 and CDRL3 of FH1 and of known VRC01-class antibodies. Tryptophan residues located exactly five amino acids away from the carboxy terminal of CDRH3 are highlighted in red. Glutamic acid residues located at position 96 in the CDRL3 of most VRC01-class antibodies are highlighted in red.

Both VH and VL antibody domains were minimally mutated with 5 amino acid changes in VH1-2*02 (H34N in CDRH1, G56D in CDRH2, I75S, S82N and L84V in FRH3) and only 2 changes in κ3-20 (S29R in CDRL1, and A43V). Overall, there was 92% amino acid homology between FH1 and glVRC01 in both their VH and VL domains. The CDRH3 of FH1 is 11 amino acids long, which is within the range of CDRH3s of VRC01-class antibodies (**Fig. 1b**) and like most VRC01-class antibodies expresses a Trp exactly five amino acids away from the carboxy terminal domain of CDRH3 (Zhou et al., 2013). The CDRL3 sequence (QHMYT) of FH1 differs from the prevalent κ3-20 CDRL3 sequence motifs (QQY/LE) present in known VRC01-class antibodies (Scheid et al., 2011; West et al., 2012; Zhou et al., 2013) (**Fig. 1b**). Thus, FH1 displays VH and VL sequence similarities to known VRC01-class antibodies.

### FH1 binds to the C1C2 gp120 domain

A 3.5 Å structure of FH1 Fab complexed with the core gp120 domain of the HxB2 Env expressed in GnTI-/- cells was generated and analyzed (**Fig 2a**). The structure revealed that FH1 binds primarily to the C1 region of the gp120 Env subunit (95% of the total buried surface area (BSA)) with some contacts in the C2 region (5% of the total BSA), both in the inner domain of gp120. FH1 binds Env with a total BSA of ∼1067 Å^2^, 62% contributed from the heavy chain (CDRH1 (8%), CDRH2 (22%), CDRH3 (28%) and the rest from FR regions) and 38% from the light chain (CDRL1 (19%) and CDRL3 (15%), the rest from FR1). In comparison, glVRC01 binds Env with a total BSA of ∼950 Å^2^, 75% contributed from the heavy chain (CDRH2 (52%), CDRH3 (11%) and the rest from FR regions) and 25 % from the light chain (CDRL1 (1%), CDRL3 (17%) and the rest from FR1).

**Figure 2.**
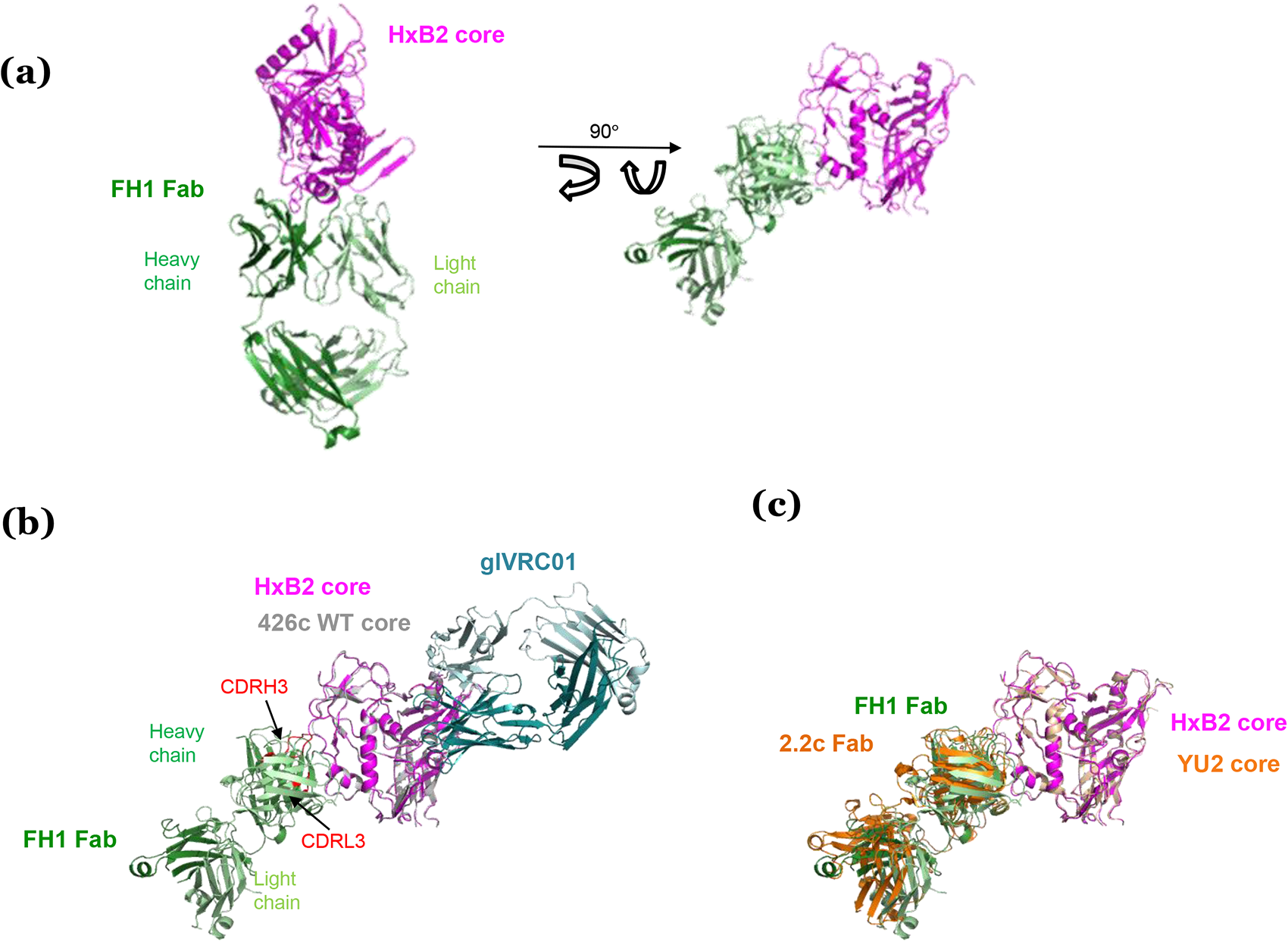
Crystal structure of FH1 Fab bound to HxB2 gp120 core indicates mode of binding. (a) Cartoon representation of the overall complex structure with HxB2 core shown in magenta and FH1 Fab shown in green (heavy chain in dark green and light chain in light green). Two views are shown. **(b)** Superposition of structure of WT 426c Core (grey) bound to glVRC01 (marine) onto that of HxB2 Core bound to FH1. The FH1 CDRH3 and CDRL3 domains are highlighted. (**c**) Superposition of YU2 Core bound to 2.2c Fab structure (orange) onto that of HxB2 Core bound to FH1 indicates a similar epitope.

Three gene-encoded amino acids in the VH1-2*02 domains of VRC01-class antibodies (Trp50_HC_, Asn58_HC_ and Arg71_HC_, Kabat numbering) play key roles in the antibody interactions with the CD4-BS of Env (highlighted in red in **Fig. 1b**). Trp50_HC_ (in FWR2) interacts with Asn280_gp120_ in Loop D, Asn58_HC_ (in FWR3) interacts with Arg456_gp120_ in V5 and Arg71_HC_ (in FWR3) interacts with Asp368 _gp120_ within the CD4 binding loop (Scharf et al., 2013; West et al., 2012). These three amino acids remain unaltered during the maturation of VRC01-class antibodies. In addition, Trp100B_HC_ in CDRH3 interacts with Asn/Asp279 in Loop D. In contrast, in the case of FH1, Trp50, Asn58 and Trp100A interact with the C1 and C2 domains, while Arg71 does not contact Env. Thus, while VRC01-class antibodies recognize the CD4-BS of Env, FH1 recognizes elements of the C1/C2 domains of gp120 (**Fig. 2b**). The overall angle of approach of FH1 to gp120, however, is similar to that of the weakly ADCC-inducing A32-like antibody, 2.2c (Acharya et al., 2014) (**Fig. 2c**).

### Env cross-recognition properties of FH1

FH1 recognizes both the 1086.C and TV1 Envs used in the HVTN100 vaccine (**Fig. 3a**). So far, only a handful of Env-derived recombinant proteins that bind germline forms of VRC01-class antibodies (‘germline-targeting’ proteins) have been designed: 426c Core, 426c OD, eOD-CT6/GT8, and BG505 GT1.1 (Briney et al., 2016; Dosenovic et al., 2015; Duan et al., 2018; Jardine et al., 2015; Lin et al., 2020; McGuire et al., 2016; Medina-Ramirez et al., 2017; Parks et al., 2019; Sok et al., 2016; Tian et al., 2016). While 426c Core expresses both the inner and outer gp120 domains of the clade C 426c Env, eOD-GT8 and 426c OD express the outer gp120 domain of the clade B HxB2 Env and 426c, respectively.

**Figure 3.**
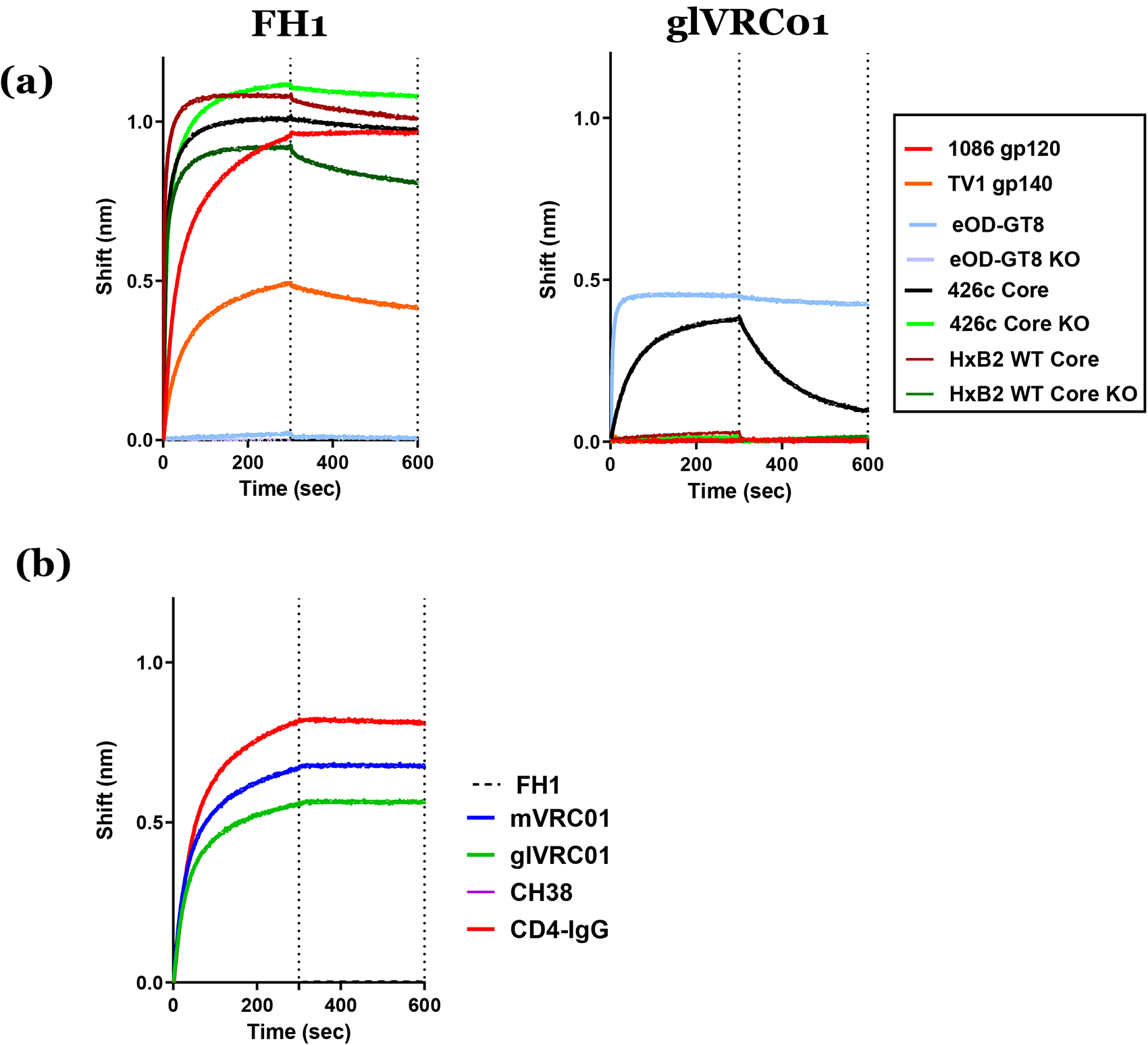
Antibody-Env interactions. (**a**) The interactions between FH1 or germline VRC01 (glVRC01) and the indicated recombinant Envs were determined by Biolayer Interferometry. KO indicates the presence of amino acid mutations that abrogate the binding of CD4-binding site antibodies (D368R, E370A, and D279A in the case of 426c Core and HxB2 Core and D368R and D279A in the case of eOD-GT8). (**b**) The binding of FH1 to eOD-GT8 monomer in the absence or presence of indicated antibodies and IgG-CD4 was monitored by Biolayer Interferometry. Mature VRC01(mVRC01), germline VRC01 (glVRC01).

FH1 did not bind eOD-GT8 (which lacks C1 and C2) (nor 426c OD, data not shown), but bound 426c Core (which expresses both C1 and C2) (**Fig. 3a**). As expected, FH1 bound equally well to a variant of 426c Core with the CD4-BS knock-out mutations (D368R, E370A and D279A) (426c Core CD4-BS KO) and recognized the HxB2 WT Core protein in a CD4-BS-independent manner (i.e., it bound the HxB2 WT Core even in the presence of the above mentioned CD4-BS KO mutations, HxB2 WT Core KO). In contrast, and as expected, glVRC01 MAb bound eOD-GT8 and 426c Core, in a CD4-BS-dependent manner, as previously discussed (Jardine et al., 2015; McGuire et al., 2016; Yacoob et al., 2016), but not the HxB2 WT Core, 1086.C or TV1 Envs.

Antibody competition experiments performed between FH1 and mVRC01, glVRC01, CD4-IgG or CH38 (an anti-C1 MAb) (**Fig. 3b**) indicated that FH1 does not compete with the anti-CD4-BS antibodies or IgG-CD4, but competes with CH38 (and with itself). Collectively, the above results confirm that FH1 recognizes a known target of vaccine-elicited antibodies with ADCC activities (Bonsignori et al., 2012; Easterhoff et al., 2017; Ferrari et al., 2011).

The affinity of FH1 to 426c Core is about 2 orders of magnitude greater than that of glVRC01 (**Fig. S2**). Not surprisingly, the affinity of FH-1 for 1086.C gp120, contained in the vaccine boost, is approximately 12-fold greater than for the 426 Core.

In summary, although the VH and VL regions of FH1 and VRC01-class antibodies share very similar amino acid sequences, the two antibodies recognize different Env domains.

### Both CDRH3 and CDRL3 dictate epitope-specificity of FH1

As discussed above, the gene-encoded portions of the VH and VL domains of glVRC01-class antibodies and those of FH1 differ in only seven amino acid positions (five in VH and two in VL). The major sequence differences are in CDRH3 and CDRL3. We therefore examined whether the differences in epitope-recognition of FH1 and glVRC01 are due to sequence variations in either CDRH3, CDRL3, or both (**Fig. 4**). For this analysis we used eOD-GT8 as the target Env as it is recognized by glVRC01, but not FH1 (**Fig. 3a**).

**Figure 4.**
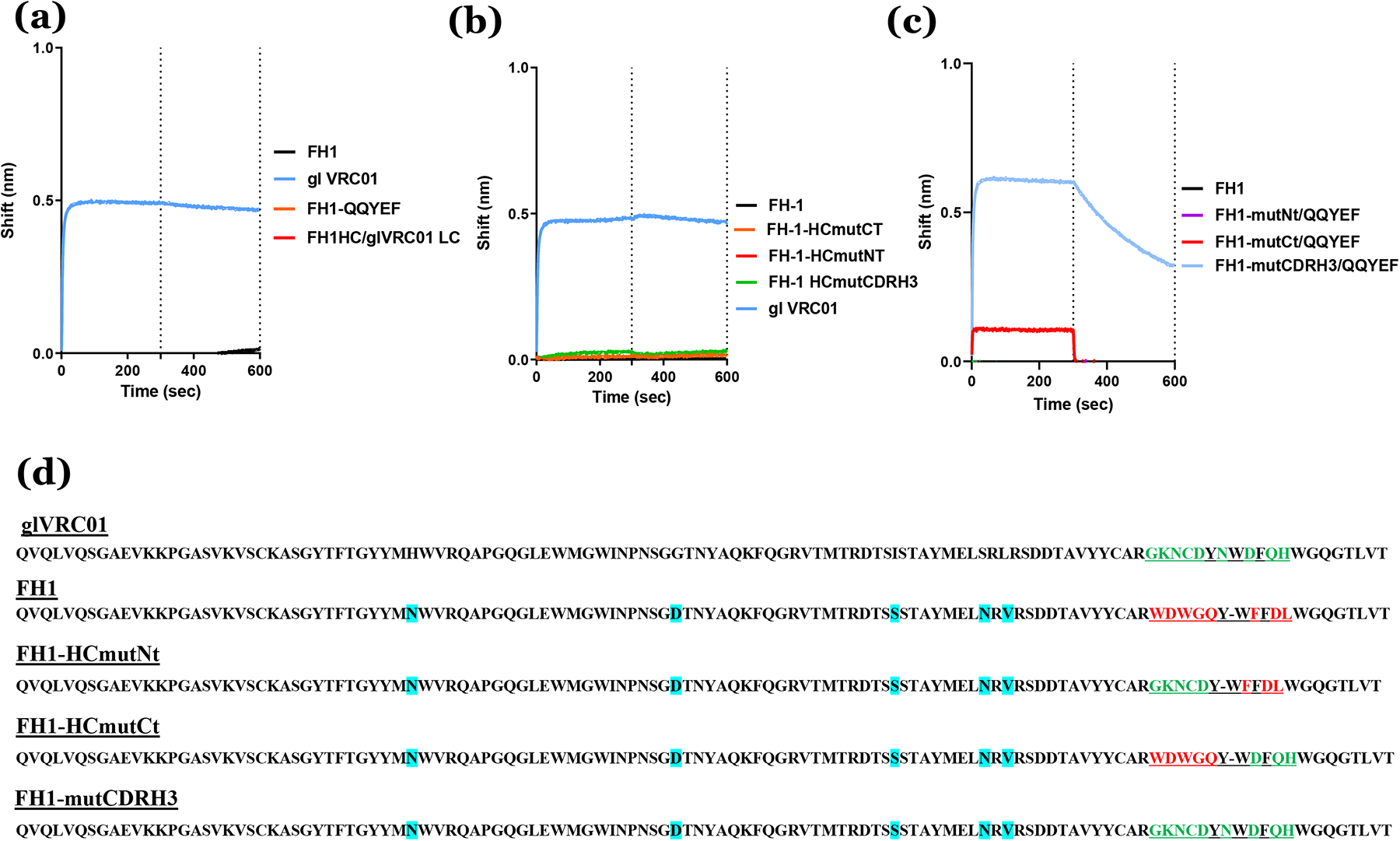
Role of CDRH3 and CDRL3 is defining the epitope specificity of FH1. The interactions between FH1 and the various FH1/VRC01 chimeric antibodies with eOD-GT8 (monomer) were determined by Biolayer interferometry. **(a)** Effect of substitutions in CDRL3, (**b**) effect of substitutions in CDRH3, and (**c**) effect of substitutions in both the CDRL3 and CDRH3. (**d**) The amino acid sequences in HC of all antibodies examined are presented. Amino acids that differ between the HC sequences of FH1 and glVRC01 are highlighted in cyan.

We first replaced the CDRL3 amino acid sequence of FH1 (QHMYT) with that of VRC01 (QQYEF) (**Fig. 4a**). That antibody, FH1-QQYEF, like FH1 did not bind eOD-GT8. We then paired the FH-1 heavy chain with the entire κ3-20 light chain of glVRC01, but that antibody (FH1 HC/glVRC01 LC) did not bind eOD-GT8 either (**Fig. 4a**).

Next, we examined whether amino acid differences in CDRH3 were responsible for the differential epitope specificities of FH1 and VRC01. To this end, we replaced the carboxy-terminal half of the CDRH3 of FH1 with that of VRC01 (FH1-HCmutCT), the amino-terminal half of the CDRH3 of FH1 with that of VRC01 (FH1-HCmutNT), or the entire CDRH3 of FH1 with that of VRC01 (FH1-HCmutCDRH3) and examined the effect of these alterations on antibody-binding to eOD-GT8. None of these antibodies bound eOD-GT8 (**Fig. 4b**). Collectively, these results indicate that the amino acid differences in the individual CDRH3 and CDRL3 domains between FH1 and VRC01 are not, by themselves, responsible for the different epitope specificities of these antibodies.

We thus examined whether the different epitope specificities were due to differences in both the CDRH3 and CDRL3 domains. To this end, we engineered chimeric antibodies expressing the HCs of the above mentioned FH1-HCmutNT, FH1-HCmutCT and FH1-HCmutCDRH3 antibodies with the LC of FH1-QQYEF and examined their binding to eOD-GT8 (**Fig. 4c**). No binding was observed with the FH1-HCmutNt/QQYEF antibody, weak binding was observed with the FH1-HCmutCt/QQYEF antibody, but robust binding was observed with the FH1-HCmutCDRH3/QQYEF antibody. We conclude, therefore, that for FH1 to change its epitope specificity from the C1/C2 domain to the CD4-BS of gp120, extensive amino acid changes in both the CDRH3 and CDRL3 domains would be required.

### FH1 has ADCC but not neutralizing potential

We evaluated the ability of FH1 to neutralize the vaccine-matched 1086c strain and the panel of 426c viral mutants that have been engineered to be sensitive to neutralization by the germline VRC01 mAb (LaBranche et al., 2018). There was no neutralization of the viruses detected at up to 50 mg/ml. We also evaluated the ability of FH1 to mediate antibody-dependent cellular cytotoxicity (ADCC) to a panel of infectious molecular clones. FH1 was able to mediate ADCC against the vaccine-matched strain 1086c, even at very low concentrations (>0.05 mg/ml), which is the same strain that was used as a gp120 to isolate this antibody (**Fig. S3**). However, FH1 was unable to mediate ADCC against any other viruses tested including TV1, which is a subtype C vaccine-matched virus, or others from a global panel of ten heterologous viruses tested.

### Competition between FH1- and VRC01-expressing B cells for Env

The results presented above, suggested that the 426c Core immunogen, which is designed to activate B cells expressing glVRC01-class Abs (McGuire et al., 2016; Parks et al., 2019), will also activate B cells expressing FH1-like B cell receptors (BCRs) which can give rise to antibodies with anti-HIV-1 ADCC activities. Potentially, if the frequency of FH1-like B cells and the relative affinity of the FH1-like BCRs for 426c Core are higher than those of glVRC01 B cells, the 426c Core may not be able to activate the latter cells. In contrast, the expectation is that the 1086 Env would only activate FH1-expressing B cells, while eOD-GT8 would activate VRC01-expressing B cells, but not FH1-expressing B cells. To investigate this directly, we engineered B cells expressing FH1 and glVRC01 BCRs and examined their activation by 1086-C4b, 426c Core-C4b and eOD-GT8-C4b self-assembling nanoparticles (7meric) at three different FH1/glVRC01 B cell ratios (10:1, 1:1 and 1:10) (**Fig. 5**). Indeed, eOD-GT8-C4b activated glVRC01 B cells but not FH1 cells irrespective of the relative FH/glVRC01 B cell ratios (**Fig. 5, top row**), 426c Core-C4b activated both B cells irrespective of the relative B cells ratios (**Fig. 5, middle row**), while 1086-C4b only activated FH1 B cells irrespective of the relative B cell ratios (**Fig 5, bottom row**). These data suggest that eOD-GT8 and 426c Core, both germline-targeting immunogens, are therefore more likely to activate glVRC01-class B cells rather than non-germline-targeting Env immunogens. Tin addition, the 426c Core may activate anti-C1C2 gp120 domain B cells that are linked with anti-HIV-1 ADCC activities.

**Figure 5.**
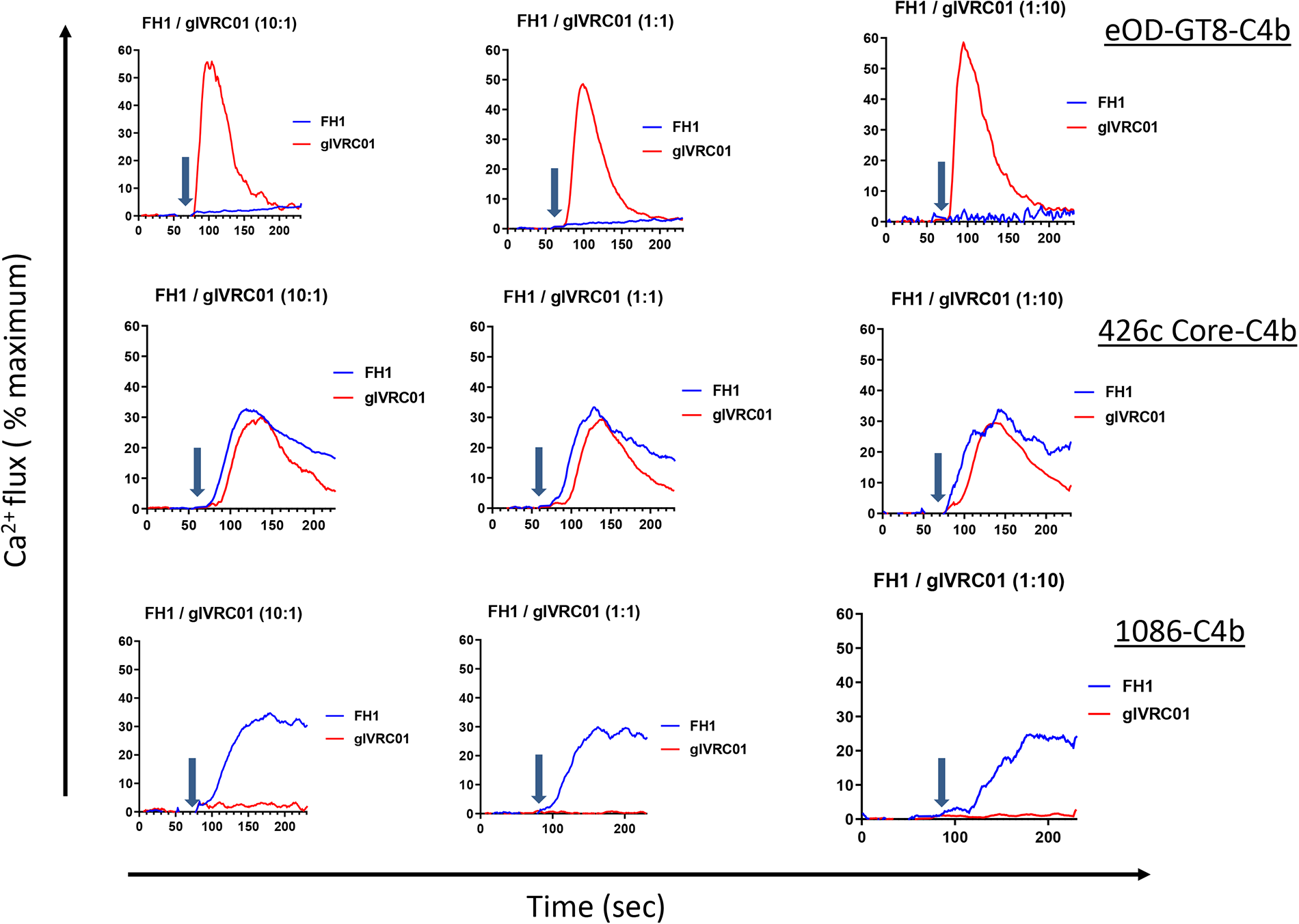
Activation of B cells expressing FH1 or glVRC01 BCRs. B cells engineered to express either FH1 or glVRC01 BCRs were mixed at the indicated ratios and incubated with eOD-GT8-C4b (top panels), 426c Core-C4b (middle panels), or 1086-C4b (bottom panels). The relative activation of FH1 (blue traces) or glVRC01 (red traces) was evaluated by monitoring changes in intracellular Ca^2+^ concentrations overtime. Arrows indicate when the indicated Envs were added.

## DISCUSSION

The VH and VL genes of FH1 are minimally mutated. This implies that the corresponding B cell did not undergo extensive somatic hypermutation and may have been activated late during the HVTN 100 immunization series. Because 1086 and TV1 were co-administered and both bind FH1, it is presently unknown which of the two immunogens initiated the activation of the corresponding naïve B cell. Still, the possibility exists that during a more prolonged germinal center reaction, the FH1 BCR may accumulate somatic mutations that would lead to a change in its epitope-specificity from the C1/C2 domain to the CD4-bs. However, here, we demonstrate that such a dramatic epitope-switch would require extensive and simultaneous changes in both the CDRH3 and CDRL3 domains. Thus, it is unlikely that FH1-like expressing B cells which become activated by non-germline-targeting Envs will become VRC01-like through the accumulation of somatic hypermutations. Our findings also support observations that despite the predominance of V gene-encoded antibody domain CDRH2 in the interaction of VRC01-class antibodies with the CD4-bs, their CDRH3 domains influence this epitope-specificity (Bonsignori et al., 2011; Yacoob et al., 2016).

Efforts to elicit VRC01-like antibodies have focused on designing recombinant Env immunogens capable of binding with high affinity to the unmutated forms of these antibodies, as expressed on naïve human B cells (Jardine et al., 2013; Jardine et al., 2016; Lin et al., 2020; McGuire et al., 2016; McGuire et al., 2013; Medina-Ramirez et al., 2017; Parks et al., 2019). Such ‘germline-targeting’ immunogens present not only the epitope of interest (the VRC01 epitope), but other epitopes recognized by non-VRC01-like antibodies, which are non-neutralizing, as well (Havenar-Daughton et al., 2018; McGuire et al., 2014a). Depending on the relative affinities and frequencies of the on-target and off-target B cells, the former B cells may not be efficiently activated by germline-targeting immunogens (Abbott et al., 2018; Dosenovic et al., 2018). Here we show that even if the off-target B cells (non-VRC01-like and non-neutralizing) are in large excess over the on-target B cells (VRC01-like), germline-targeting immunogens can still activate the on-target B cells. In contrast, even if the on-target B cells are in large excess over the off-target B cells, a non-germline-targeting immunogen will not activate the former cells but will readily activate the latter cells.

The observation that FH1 and glVRC01-like antibodies display high degrees of amino acid sequence homology, but different epitope specificities, highlights the importance of combining paired VH/VL gene sequence analysis with actual antibody-binding and structural analysis to properly assess the success of upcoming human immunizations aiming at activating VRC01-class BCRs.

In summary our study provides direct evidence that Env immunogens that are not designed to engage the unmutated forms of VRC01-class antibodies are unlikely to activate the corresponding B cells in humans. Specifically, our study supports the proposal that the *in vivo* activation of B cells expressing the precursor BCR forms of VRC01-class bnAbs will require specifically designed protein immunogens.

## ACKNOWLEDGEMENTS

This study was funded by NIH grants R01AI081625 (LS), R01AI104384 (LS), UM1AI144462 (subaward MJM) and UM1AI068618 (MJM). We appreciate the efforts of the HVTN 100 Protocol Team, including protocol chairs, Drs. Linda Gail Bekker and Fatima Laher, and the study participants in the conduct of that trial. We thank the Fred Hutchinson Cancer Research Center’s Vector Core for generating AAV viruses.

## MATERIALS AND METHODS

### HVTN 100 clinical trial

HVTN 100 was a randomized, controlled, double-blind study phase 1-2 clinical trial, which enrolled 252 healthy HIV-uninfected 18- to 40-year-old participants at 6 sites in South Africa. The relevant research ethics committees approved the study. All participants gave written informed consent in English or their local language. Additional detail about the HVTN 100 study design, eligibility criteria, participants and their baseline characteristics, randomization, blinding, study products is available in previous reports (Bekker et al., 2018). The trial was registered with the South African National Clinical Trials Registry (DOH-27-0215-4796) and ClinicalTrials.gov (NCT02404311).

### Recombinant Envelope protein and antibody expression and purification

Details on the expression and purification of recombinant HIV-1 envelope proteins and of monoclonal antibodies have been extensively presented (Lin et al., 2020; Parks et al., 2019; Sellhorn et al., 2009; Sellhorn et al., 2012). The Envs employed during the Ca2+ flux assays were multimerized using the C4b-binding protein oligomerization motif, as we previously described in detail (Lin et al., 2020; McGuire et al., 2016; Parks et al., 2019).

### Complex Crystallization and X-ray data collection

HxB2 was transfected in GNTI-/- cells and purified with GNA lectin and size exclusion chromatography (SEC). FH1 Fab was obtained by digesting FH1 IgG with Lys C, collecting the Flow Through from a Protein A column and purifying it by SEC. HxB2 and FH1 Fab were incubated at a 1.1 molar excess of Fab, treated with EndoH for 1 hr at room temp and the complex was purified over SEC. The complex was screened against the Hampton Crystal HT, ProPlex HT-96, and Wizard Precipitant Synergy #2 crystallization screens. The NT8 robotic system (FORMULATRIX^®^) was used to set initial sitting drop crystallization trials. Following initial hits, crystallization conditions were optimized using hanging drop vapor diffusion. Crystals were grown in 0.1 M NH4SO4, 20% PEG 1500, 0.1M Tris pH 7.5 at 10 mg/ml. Crystals were flash frozen in 0.15M (NH4)_2_SO4, 30% PEG 1500, 0.15M Tris pH 7.5) supplemented with 20% Ethylene Glycol. Data sets were processed using HKL2000 (Otwinowski and Minor, 1997), and initial models were generated using molecular replacement in Phenix (Adams et al., 2010). Following molecular replacement, iterative model building and refinement was achieved using COOT (Emsley and Cowtan, 2004) and Phenix, respectively.

### Biolayer Interferometry (BLI)

BLI experiments were performed as previously described (Lin et al., 2020; Parks et al., 2019). Briefly, kinetic analyses were performed using recombinant Fabs loaded onto FAB2G biosensors (ForteBio, Cat#: 18-5126) (@ 40 ug/ in 1XPBS) and 2-fold dilutions of Env monomers. The assay parameters are the same as for measuring the binding of IgG, but with an extended dissociation phase of 600 s. Curve fitting used to determine relative apparent antibody affinities for envelope was performed using a 1:1 binding model and the data analysis software (ForteBio). Mean kon, koff, and KD values were determined by averaging all binding curves within a dilution series having R2 values of greater than 95% confidence level.

### HIV-1 neutralization and ADCC assays

All neutralization experiments were performed as previously described (LaBranche et al., 2018). ADCC was determined by a luciferase (Luc)-based assay as previously described (Chua et al., 2021; Liao et al., 2013). Briefly, CEM.NKR_CCR5_ cells (NIH AIDS Reagent Program, Division of AIDS, NIAID, NIH from Dr. Alexandra Trkola) (PMC112928) were used as targets after infection with the HIV-1 IMCs. Recombinant FH1 mAb was tested across a range of concentrations using 5-fold serial dilutions starting at 50 µg/mL. The final read-out was the luminescence intensity (relative light units, RLU) generated by the presence of residual intact target cells that have not been lysed by the effector population in the presence of ADCC-mediating mAb (ViviRen substrate, Promega, Madison, WI). The % of specific killing was calculated using the formula: percent specific killing = [(number of RLU of target and effector well − number of RLU of test well)/number of RLU of target and effector well] ×100. In this analysis, the RLU of the target plus effector wells represents spontaneous lysis in absence of any source of Ab. The ADCC positive cutoff was 15% specific killing based on our previous reports. The mAb palivizumab (Synagis), which mediates ADCC (Hiatt et al., 2014) but is specific for respiratory syncytial virus, and a cocktail of HIV-1 mAbs (HIV-1 mAb mix) demonstrated to mediate ADCC [A32 (Ferrari et al., 2011), 2G12 (Trkola et al., 1996), CH44 (Moody et al., 2015), and 7B2 (Sadraeian et al., 2017)] were used as negative and positive controls, respectively.

### Generation of FH1- and glVRC01-A20 cell lines

FH1- and glVRC01-A20 cell lines were generated using CRISPR technique as described previously (Moffett et al., 2019). The sgRNA muIgH367 sequence is TTATACAGTATCCGATGCAT**AGG (**PAM site in bold) targeting the region between the last JH gene and the Eu enhancer which is upstream of endogenous IgH constant region. 18 umol Cas9 proteins (Invitrogen) and 54 umol sgRNA (Synthego) were precomplexed in 5 ul Neon Buffer R at room temperature for 20 minutes. 250, 000 A20 cells were resuspended in 7ul R buffer, added to the Cas9/sgRNA complex and electroporated using 20-ms pulse at 1725 V using the Neon Transfection System (Invitrogen). The cleavage efficiency of sgRNA was about 60% as assessed by Synthego ICE CRISPR software after PCR amplification and sequencing of the targeted genomic DNA. After electroporation, cells were recovered in pre-warmed RPMI complete media without antibiotics for 30 minutes, and then transduced by FH1 or glVRC01 AAV viruses. After expanding for 3 days, cells were stained WITH Q3W24 BGF BITH BUV395 anti-CD19 (BD Bioscience), AF647 conjugated 426cDMRScore-c4b and Near-IR live dead dye (ThermoFisher Scientific) to sort out the live, CD19 positive, Env binding population on FACSAria (BD Biosciences). AAV viruses were produced by co-transfection of FH1- or glVRC01-AAV plasmids, serotype 6 capsid, and adenoviral helper plasmids into HEK293 cells using polyethylenimine. Viruses were purified over iodixanol gradient. Their titers were approximately 1x10^9^ per microliter. FH1 and glVRC01 AAV plasmids were constructed by In-Fusion (Takara) recombination of ECoRV linearized AAV backbone plasmid PCH19-mb3AAV-HR (gift from Dr. Justin Taylor’s lab) and FH1 or glVRC01 gBlocks (IDT). The FH1 and glVRC01 gBlocks contain the full-length light chain, the 57-amino acid glycine-serine linker, and the heavy chain VDJ, plus In-Fusion homology sequences to the backbone plasmid.

### Calcium influx

Calcium influx experiments were performed as previously described (McGuire et al., 2014a). Briefly, FH1- or glVRC01-expressing A20 cells were resuspended in RPMI complete media (1.5x10^6^/ml), mixed (1:1 in volume) with Fluo 4 dye (Fluo 4 Direct Calcium Assay Kit, ThermoFisher Scientific) and incubated at 37°C for 30 minutes. Fluorescence was recorded for 1 minute (baseline) and 0.25uM of Env was added and the fluorescence was recorded for 3 minutes, at which point ionomycin was added (2ug/ml). The Env proteins tested were eODGT8-c4b, 426cDMRSCore-c4b and 1086-C4b. Data were analyzed using the Kinetics function in FlowJo software. Baseline fluorescence was subtracted from fluorescence intensity at all time points using Prism 7 software. Calcium influx in mixed FH1- and glVRC01-expressing A20 cells were performed as previously described (McGuire et al., 2014a). Briefly, FH1- and glVRC01-A20 cells (1.0x10^6^) were labeled with 1uM CellTrace Violet and 1uM CellTrace Yellow (ThermoFisher Scientific) respectively, according to manufacturer’s protocol. The cells were washed, resuspended in RPMI complete media and stained with Fluo 4 dye. FH1- and glVRC01-expressing A20 cells were then mixed at 1:10, 1:1 or 10:1 ratios and incubated with eODGT8-c4b, 426cDMRScore-c4b or 1086-c4b as described above. Analysis was performed In FlowJo, FH1 and glVRC01 cells were gated on CellTrace Violet and CellTrace Yellow positive separately and graphed for Fluo 4 fluorescence intensity over time.

---

**Supplemental Figure S1. (a)** Timeline of vaccine administration in HVTN 100. Numbers along the timeline indicate months since enrollment. Blue syringes denote administration of ALVAC, while red syringes denote administration of the bivalent rec gp120 mix (1086 and TV1). The red arrow indicates the visit (2 weeks post-5^th^ immunization at visit 12) during which Env+ BCR repertoire analysis was performed and FH1 was isolated.

**Supplemental Figure S2. Kinetic analysis** Kinetic analyses were performed by BLI using recombinant Fabs loaded onto FAB2G biosensors and 2-fold dilutions of the indicated Env monomers.

**Supplemental Figure S3. ADCC activities** The ADCC activities of FH1, a cocktail of anti-HIV-1 mAbs (A32, 2G12, CH44 and 7B2) and of mAb palivizumab (Synagis) were determined against the indicated viruses. The ADCC positive cutoff was 15% specific killing based on our previous reports.

