## Supplemental Figures for "Characterization of a vaccine-elicited human antibody with sequence homology to VRC01-class antibodies that binds the C1C2 gp120 domain"

#### Figure S1

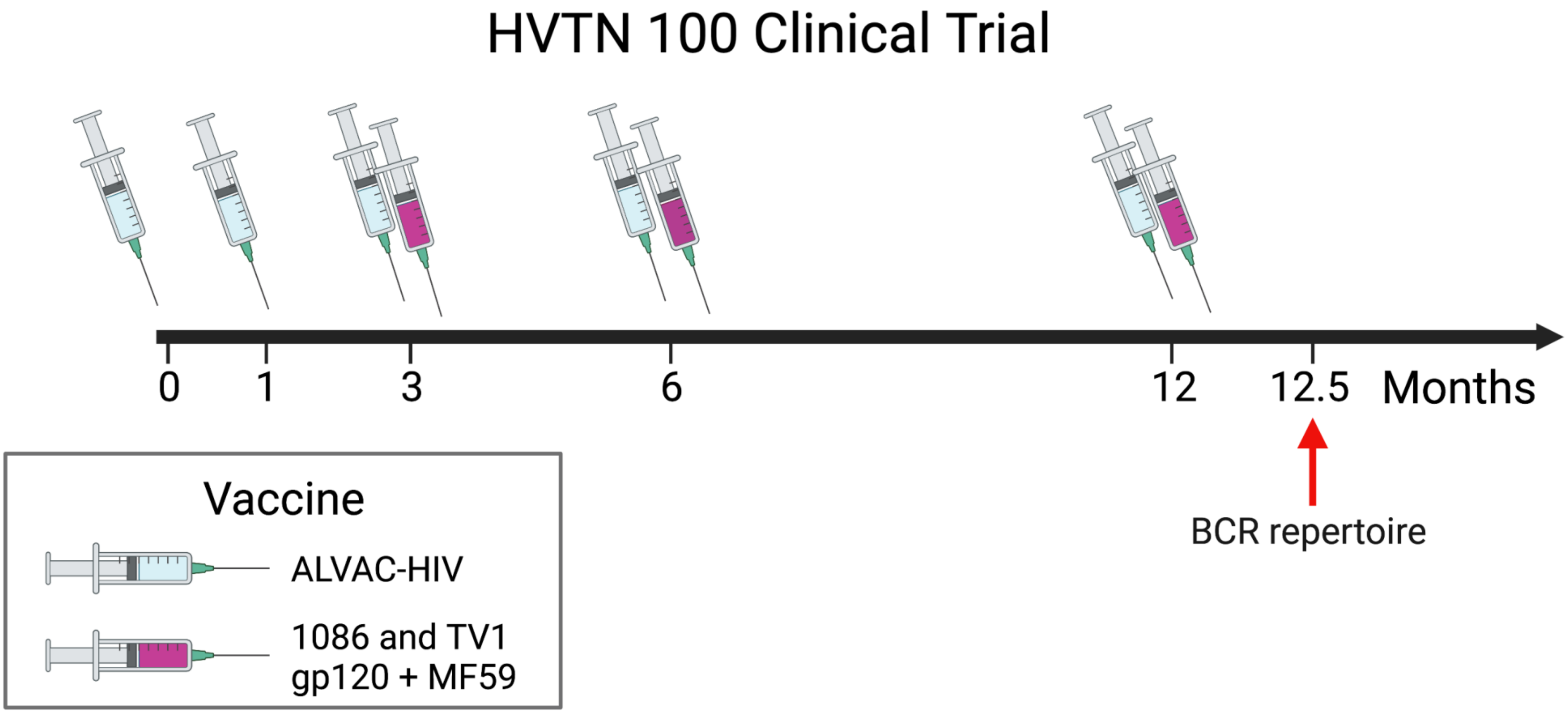
HVTN 100 Clinical Trial
0
1
3
6
12
12.5
Months
BCR repertoire
Vaccine
ALVAC-HIV
1086 and TV1
gp120 + MF59

### Figure S2

| | | KD (M) | KD Error | kon(1/Ms) | kon Error | kdis(1/s) | kdis Error | Full R^2 |
|---|---|---|---|---|---|---|---|---|
| FH-1 Fab | DMRScore gp120 | 2.64E-09 | 8.62E-11 | 2.43E+05 | 2.01E+03 | 2.79E-05 | 4.64E-06 | 0.9865 |
| | 1086 gp120 | 2.14E-10 | 2.71E-11 | 1.63E+05 | 2.23E+03 | 2.06E-05 | 3.05E-06 | 0.9904 |
| | eOD-GT8 | - | - | - | - | - | - | - |
| gl VRC01 Fab (3-20) | DMRScore gp120 | 6.09E-07 | 9.44E-09 | 9.67E+03 | 1.50E+02 | 2.48E-03 | 1.13E-05 | 0.9878 |
| | 1086 gp120 | - | - | - | - | - | - | - |
| | eOD-GT8 | 1.67E-09 | 5.48E-11 | 1.44E+05 | 1.51E+03 | 1.57E-04 | 4.04E-06 | 0.9715 |

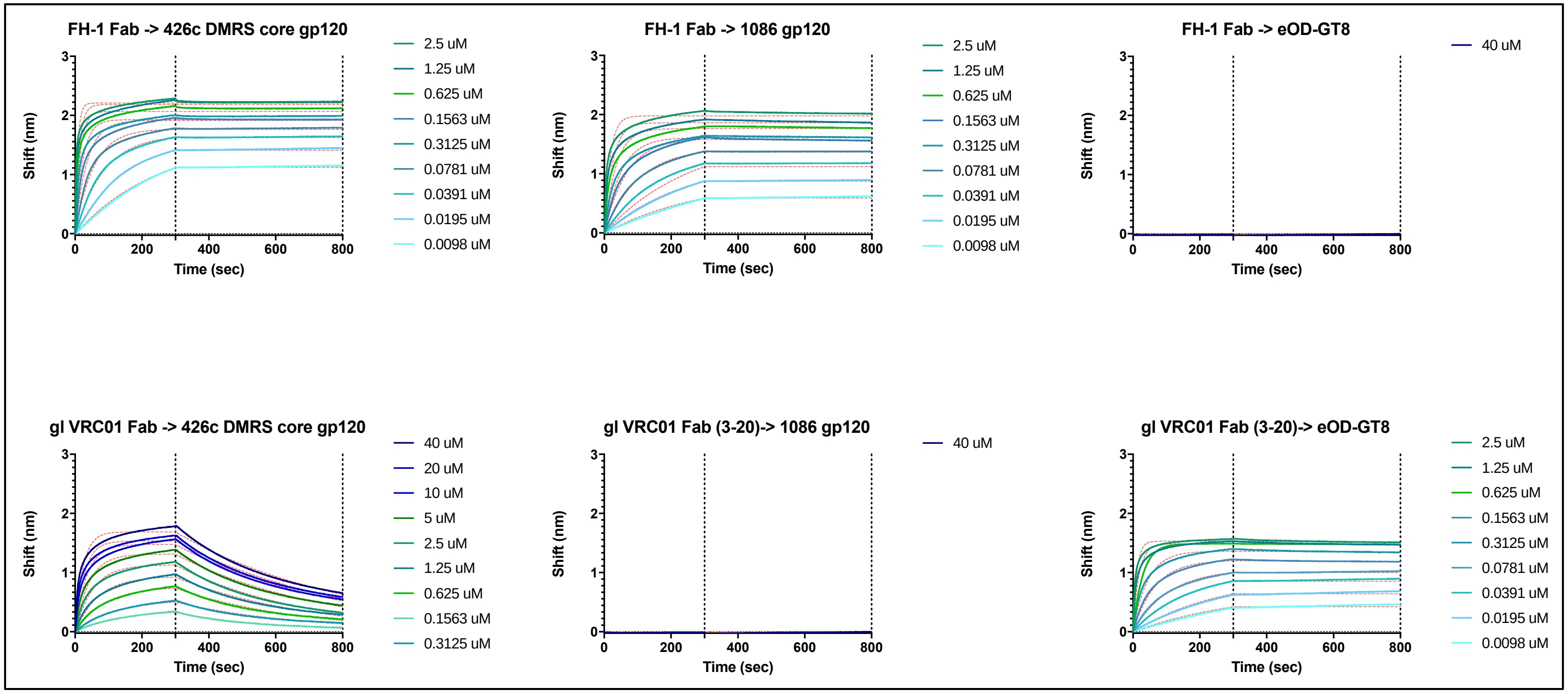

### Figure S3

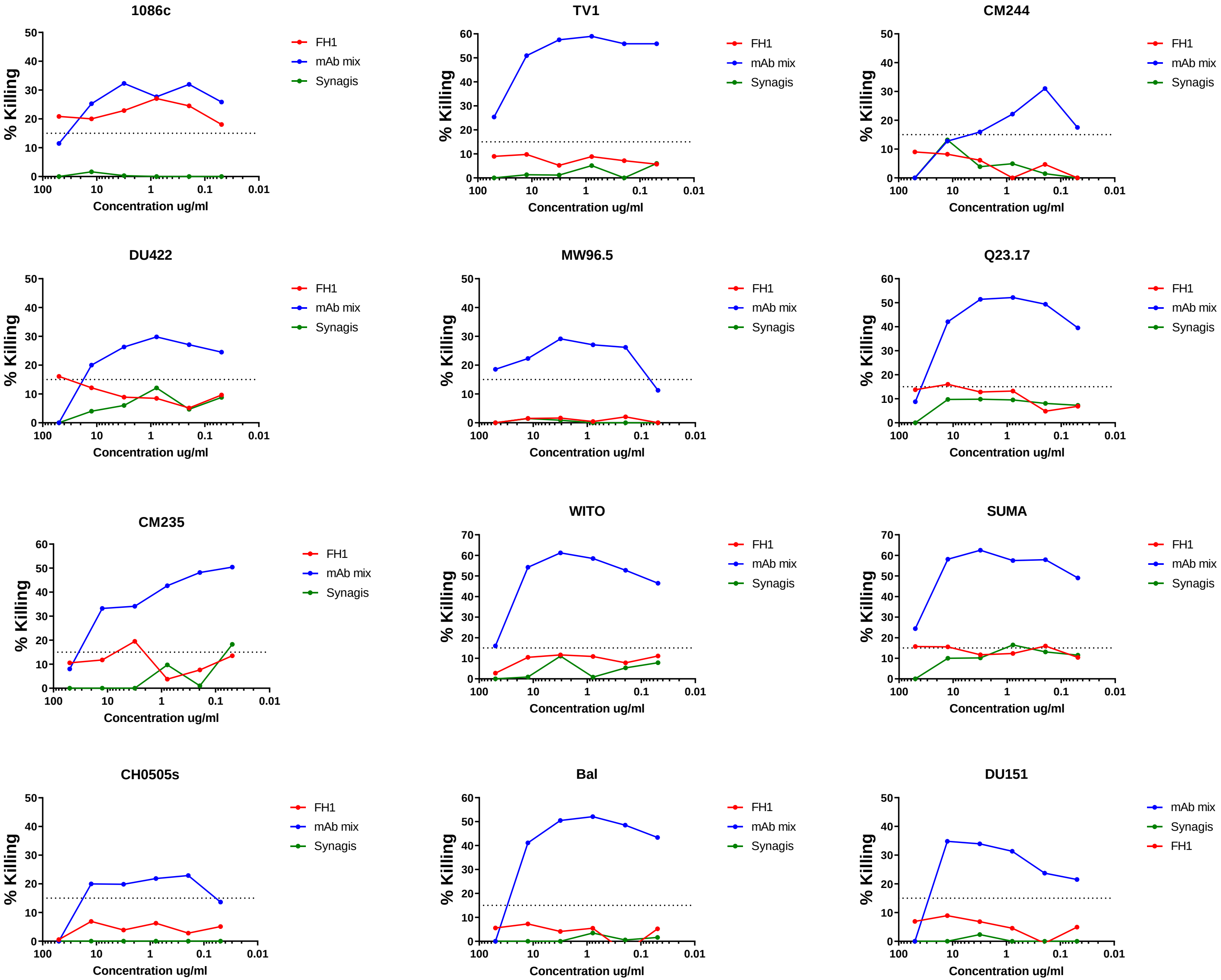
1086c
TV1
CM244
DU422
MW96.5
Q23.17
CM235
WITO
SUMA
CH0505s
Bal
DU151
% Killing
Concentration ug/ml
100
10
1
0.1
0.01
FH1
mAb mix
Synagis
